# Interpreting Lung Cancer Health Disparity at Transcriptome Level

**DOI:** 10.1101/2025.01.09.632292

**Authors:** Masrur Sobhan, Md Mezbahul Islam, Ananda Mohan Mondal

## Abstract

Lung cancer is a leading cause of cancer-related mortality, with disparities in incidence and outcomes observed across different racial and sex groups. Understanding the genetic factors of these disparities is critical for developing targeted treatment therapies. This study aims to identify both patient-specific and cohort-specific biomarker genes that contribute to lung cancer health disparities among African American males (AAMs), European American males (EAMs), African American females (AAFs), and European American females (EAFs). The real-world data is highly imbalanced with respect to race, and the lung cancer dataset is no exception. So, classification with race labels will generate highly biased results toward the larger cohort. We developed a computational framework by designing the classification problems with disease conditions instead of races and leveraging the local interpretability of explainable AI, SHAP (SHapley Additive exPlanations). This study used three disease conditions of lung cancer, including Lung Adenocarcinoma (LUAD), Lung Squamous Cell Carcinoma (LUSC), and Healthy samples (HEALTHY) to design four classification tasks: one 3-class problem (LUAD-LUSC-HEALTHY) and three 2-class problems (LUAD-LUSC, LUAD-HEALTHY, and LUSC-HEALTHY). This multiple-classification approach allows a LUAD patient to be interrogated via three classification problems, namely LUAD-LUSC-HEALTHY, LUAD-LUSC, and LUAD-HEALTHY, thus providing a robust approach of retrieving disparity information for individual patients through the local interpretation of SHAP.

The proposed method successfully discovered the sets of genes and pathways related to health disparities in lung cancer between two cohorts, including AAMs vs. EAMs, AAFs vs. EAFs, AAMs vs. AAFs, and EAMs vs. EAFs. The discovered list of genes and pathways provide a short list for biological scientists to conduct wet lab experiment.

## I. Introduction

In the United States, lung cancer is the leading cause of death. In 2023, approximately 350 deaths per day occurred from lung cancer [1], 81% of which was caused by cigarette smoking directly, with an additional 3% due to second-hand smoke [2]. The estimated new cases (75 vs. 67 in 100,000) and death rates (51 vs. 45 in 100,000) in lung cancer are *higher* among African American Males (AAMs) than in European American Males (EAMs) [1]. On the other hand, the estimated new cases (47 vs. 56 in 100,000) and death rates (28 vs. 33 in 100,000) are *lower* among African American Females (AAFs) than in European American Females (EAFs) [1]. Thus, there exists a complex disparity puzzle in lung cancer etiology between African Americans (AAs) and European Americans (EAs) in terms of both race and sex, more specifically, the *disparity between AAMs vs. EAMs and AAFs vs. EAFs*.

Cigarette smoking is considered the strongest risk factor for lung cancer, but smoking alone cannot explain the disparity of lung cancer development between AAs and EAs [3]. Based on genome-wide association studies (GWAS) for 13 cancers, Sampson *et al*. [4] found that only 24% of lung cancer’s heritability can be attributed to genetic determinants of smoking, which indicates the complex nature of heterogeneity exists in lung cancer development leading to health disparity. In a recent differential gene expression analysis (DGEA) using mRNA and miRNA expression profiles between lung tumors and normal adjacent to tumors (NAT) of AAs and EAs, researchers discovered 3,500 differentially expressed probes from AAs and 4,707 differentially expressed probes from EAs [3]. Many probes were common, along with 637 AA-specific and 1,844 EA-specific probes. Surprisingly, principal component analysis (PCA) showed that AA-specific differentially expressed probes could separate lung tumors and NAT samples in both AAs and EAs. This observation suggests that AA-specific differentially expressed probes/genes discovered using DGEA analysis cannot be considered AA-specific risk factors. The recent genome-wide association studies (GWAS), considering a large cohort of cases and controls for African Americans (AAs) [5] and European Americans (EAs) [6], failed to discover AA-specific susceptible loci since both studies discovered the same two loci near plausible candidate genes, *CHRNA5* and *TERT*, on 15q25 and 5p15 respectively, associated with lung cancer risk in both AAs and EAs.

Feature selection has long been a valuable approach for identifying biomarker genes in various cancers [7], [8]. Several studies have focused on discovering lung cancer biomarkers. For example, Sobhan et al. employed a deep learning-based feature selection algorithm to identify key genes that can differentiate lung cancers between AAMs and EAMs [9]. Additionally, researchers have explored the use of explainable machine learning techniques, such as SHAP (**SH**apley **A**dditive Ex**P**lanations) [10], to identify patient-specific biomarker genes in lung cancer patients [11]. While this research effectively identified patient-specific biomarkers for two types of lung cancer, including Lung Adenocarcinoma (LUAD) and Lung Squamous Cell Carcinoma (LUSC) patients, it did not address the race and sex-specific lung cancer health disparity.

It is clear from the existing literature mentioned above that no study considered lung cancer disparity between AAMs vs. EAMs and AAFs vs. EAFs. Most studies are based on AAs and EAs, meaning the cohort is a male and female mix. DGEA and GWAS’s inability to discover AA-specific risk loci is because these approaches are cohort-based and largely ignore the genetic and epigenetic variability of individuals or intratumor heterogeneity (ITH), and result in population-based conclusions or one-size-fits-all solution [12]. A recent committee on “Using Population Descriptors in Genetics and Genomics Research” concluded that there is no one-size-fits-all solution since research conducted using genomics data is broad and varied [13]. Thus, there is an overarching need for an alternative approach to discover risk factors that *<u>can</u> <u>explain</u>* the lung cancer health disparity not only between AAs and EAs but also between AAMs and EAMs, and AAFs and EAFs. To overcome the shortcomings of existing cohort-based approaches, we developed a computational framework to discover and interpret the lung cancer health disparity by leveraging machine learning and the explainable AI approach, SHAP [10]. The ***salient features and contributions*** of this study are enumerated below.

- We *<u>assume</u>* that the local interpretation of SHAP or interpretation of each patient of AA and EA cohorts under different disease classes/labels such as LUAD, LUSC, and HEALTHY will help extract patient-specific significant genes reflecting patient-specific disparity related to those disease conditions.
- We *<u>also</u> <u>assume</u>* that interpretation of a patient using the different combinations of disease classes (LUAD-LUSC-HEALTHY; LUAD-LUSC; LUAD-HEALTHY; LUSC-HEALTHY) would help derive a robust set of patient-specific genes. Each of the LUAD patients (irrespective of race and sex) was interrogated via three classification problems containing LUAD cohort, namely LUAD-LUSC-HEALTHY, LUAD-LUSC, and LUAD-HEALTHY. In similar way, each LUSC patient was interrogated via three classification problems containing LUSC cohort (LUAD-LUSC-HEALTHY, LUAD-LUSC, and LUSC-HEALTHY).
- We explored SHAP in a *<u>bottom-up</u> <u>approach</u>* (going from patient-specific biomarker genes to cohort-specific biomarker genes) to decipher the disparity between any two cohorts of patients, including AAMs vs. EAMs, AAFs vs. EAFs, AAMs vs. AAFs, and EAMs vs. EAFs.
- Note that the classification problems are designed based on disease conditions (i.e., LUAD, LUSC, and HEALTHY) to avoid the race and sex-specific imbalance in the dataset, which is innovative in discovering the health disparity. The data is highly imbalanced regarding race (AA:EA = 75:576). But input to the SHAP is a classification problem, which will be set up with the disease conditions LUAD (n = 356), LUSC (n = 295), and Healthy (n = 313) as class labels, meaning the data is balanced (each class consists of ~300 samples), as shown in Table I.
- Since transcriptome mirrors both genomic and epigenomic variability or ITH [14], this study interrogated individual patients via *<u>classification problems</u> <u>designed</u> <u>using</u> <u>transcriptome</u>* or expression profiles of ~20,000 genes.

**TABLE I.** Data Distribution Of LUAD, LUSC, And Healthy Samples. AA and EA represents African American and European American respectively.

| DISEASE | TCGA TUMOR |  | TOTAL |
| --- | --- | --- | --- |
|  | AA | EA |  |
| LUAD |  |  |  |
| MALE | 20 | 136 | 156 |
| FEMALE | 27 | 173 | 200 |
| LUSC |  |  |  |
| MALE | 16 | 195 | 211 |
| FEMALE | 12 | 72 | 84 |
| GTEx NORMAL SAMPLES |  |  | 313 |
| TOTAL |  |  | 964 |

## II. Materials and Methods

### A. Data Collection

RNA-seq gene expression data for LUAD and LUSC was downloaded from the publicly available MSKCC GitHub repository [15]. The analysis utilized normalized FPKM (Fragments Per Kilobase of transcript per Million mapped reads) gene expression data. In addition, gene expression data from healthy tissue samples were also downloaded from the same source. The disease samples were originally sourced from The Cancer Genome Atlas (TCGA) [16], while the healthy control samples were from the Genotype-Tissue Expression (GTEx) project [17]. Typically, RNA-seq data from different studies are not directly comparable due to differences in sample processing, data normalization, and potential batch effects. However, the MSKCC repository provides pre-processed data where such biases have been corrected, enabling reliable comparative analysis across the TCGA and GTEx datasets.

### B. Data Preparation

The initial dataset comprised 503 LUAD, 489 LUSC, and 423 healthy samples, totaling 1,415 samples. After removing duplicate entries by retaining only the first instance of each duplicate ID, the dataset was reduced to 1,401 samples. The healthy cohort included samples from GTEx healthy tissues and normal adjacent to tumor (NAT) samples from LUAD and LUSC cases. However, for this study, NAT samples were excluded, resulting in 313 healthy samples. To further refine the dataset, we included only samples belonging to the ‘non-Hispanic or Latino’ ethnic group, and excluded any samples lacking race or sex information. This filtering process yielded a final dataset of 964 samples, as shown in Table I with race and sex-specific breakdown. Note that the GTEex data do not have race and sex information, and it is not an issue for this study since the classification problems were designed based on disease conditions (LUAD, LUSC, and HEALTHY). The gene expression data consisted of 19,648 genes or features, which were utilized for subsequent analysis. The final dataset, as shown in Table I, was used to classify three classes-LUAD, LUSC, and HEALTHY using various machine learning algorithms.

### C. Study Flow Diagram

The overall pipeline of this study is shown in Fig. 1. The objective of this study is to delineate the race and sex-specific health disparities between two cohorts (AAMs vs. EAMs, AAFs vs. EAFs, AAMs vs. AAFs, and EAMs vs. EAFs) in two types of lung cancers (LUAD and LUSC) leveraging machine learning algorithms and the explainable AI, SHAP.

**Fig. 1.**
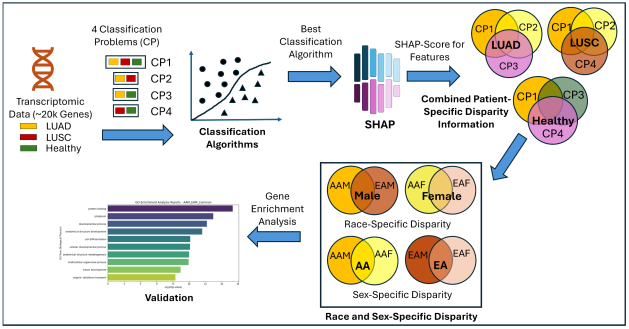
Overall flow diagram to analyze race and sex-specific health disparity in lung cancer patients.

### D. Machine Learning Algorithms

Six classification algorithms were employed in this study. These includes a 2D Convolutional Neural Network (2D-CNN) as utilized in [11], representing deep learning approach; logistic regression (LR), a regression-based method; naïve Bayesian classifier (NB), a probabilistic model; support vector machine (SVM), a kernel-based method; and two tree-based methods, random forest (RF) and extreme gradient boosting (XGBoost). A five-fold cross-validation procedure was conducted to evaluate the performance of these classifiers.

### E. Classification Problems

In this study, we formulated four distinct classification problems using three class labels, including LUAD, LUSC, and HEALTHY. One 3-class problem (LUAD-LUSC-HEALTH) and three 2-class problems (LUAD-LUSC, LUAD-HEALTHY, and LUSC-HEALTHY). The rationale behind this approach is that each classification problem may highlight different aspects of heterogeneity in lung cancer, thereby providing a more comprehensive understanding of lung cancer disparity. By implementing multiple classification tasks that focus on different pairs or groups of classes, we can identify distinct feature importance profiles that might not be apparent in a single classification task. This strategy enhances our understanding of the unique and overlapping biomarkers across classes, which is particularly valuable for distinguishing the specific characteristics of LUAD, LUSC, and HEALTHY tissues.

### F. Hyperparameter tuning

Initially, five machine learning algorithms were implemented using their default hyperparameters (HPs), while the 2D-CNN model was applied with the HPs specified in [11] for classifying LUAD, LUSC, and healthy samples. Among these, XGBoost was the best-performing classifier. Next, we tuned the HPs for the XGBoost model to enhance model accuracy for classifying LUAD, LUSC, and Healthy. The range of hyperparameters and the best values are shown in Table II. Although this approach yielded slight improvement in accuracies, it was highly time-consuming and computationally intensive to tune the HPs. Therefore, we decided to proceed with the results obtained using XGBoost with default hyperparameter values for all the classification problems. Additionally, we ran the programs ten times, each with a different seed for random state ranging between 10 and 100 in intervals of 10. The experiment with different random states was conducted because the previous study [18] found that the model produced slightly different accuracies at different random states. But in this study, the XGBoost algorithm consistently produced the same result, demonstrating the robustness of the findings in this study.

**TABLE II.**
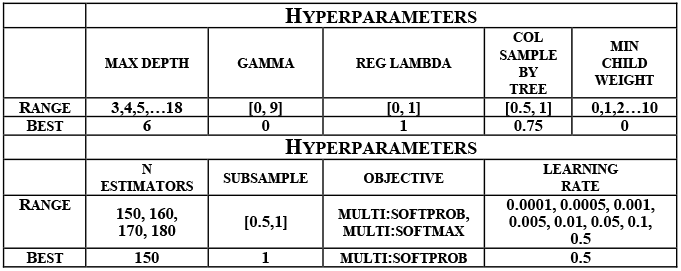
Ranges of hyperparameters and the best hyperparameters for XGBoost classifier.

### G. Local Feature Interpretation using SHAP

The SHAP is a powerful XAI tool based on game theory that helps us understand how machine learning models make decisions. SHAP assigns a score to each feature for every sample, showing how much each feature affects the model’s prediction. To calculate SHAP scores, it takes the machine learning model and the samples as input to observe how a specific feature changes the model’s prediction. It does this by comparing the model’s output with and without the feature of interest, across different combinations of features, called coalition sets. The differences in predictions are calculated for each of these sets. By averaging these differences across all possible combinations, SHAP score is calculated for a feature, which tells us how important that feature is for a particular prediction, also known as local feature interpretation.

In this study, SHAP was used to generate scores for all 964 samples across LUAD, LUSC, and healthy groups, encompassing 19,648 genes or features. A higher SHAP score indicates greater importance of a feature. Subsequently, the genes for each patient were ranked in descending order based on their SHAP scores. The top 100 genes from this ranked list were selected, as these genes are believed to contain critical risk information for the patient and are therefore referred to as patient-specific biomarker genes that carries patient-specific disparity information.

### H. Combined Patient-specific biomarker genes

Each LUAD sample was analyzed across three distinct classification tasks: LUAD-LUSC-HEALTHY, LUAD-LUSC, and LUAD-HEALTHY. We only considered the common correctly predicted samples from these three classification problems. As a result, each sample yielded three different sets of patient-specific biomarker genes, corresponding to each classification. Next, by taking the union of these three gene sets, we obtain a comprehensive list of patient-specific biomarker genes that encapsulate the combined disparity information across all classification tasks.

The same procedures were applied to LUSC and HEALTHY samples.

### I. Cohort-specific disparity information

To determine the disparity information for the African American (AAM) cohort, we performed a union operation on the lists of combined patient-specific disparity-related genes within this cohort. By aggregating these gene sets, we capture a wide spectrum of genetic markers that contribute to disparities in AAMs. This approach allows us to identify and analyze the unique biomarker patterns to identify various disease and risk factors in the AAM cohort by providing insights into cohort-specific health disparities. The same approaches were conducted to identify the disparity-related genes in AAF, EAM, and EAF cohorts.

### J. Disparity among two cohorts

By comparing cohort-specific disparity information of two cohorts, we obtained three sets of genes: one set is common between two cohorts, and the other two sets are cohort-specific. The cohort-specific set of genes are considered the cohort-specific biomarker genes.

## III. Results

### A. Selecting the Best Machine Learning Approaches

Table III shows the accuracies of six machine learning algorithms (CNN, LR, NB, SVM, RF, and XGBoost) in four classification problems. The XGBoost algorithm performs better than other five algorithms with the default values of corresponding hyperparameters. Then we used hyperparameter tuning for XGBoost and the improvement with tuned parameter was not significant, as shown in the last column. So, for the further analysis of interpretation, we used XGBoost with default values.

**TABLE III.** Acuracies of Machine Learning Algorithms in Four Classification Problems.

| Classification Problems | Using Default Values of Hyperparameters |  |  |  |  |  | Tuned |
| --- | --- | --- | --- | --- | --- | --- | --- |
|  | CNN | LR | NB | SVM | RF | XGBoost | XGBoost |
| LUAD-LUSC-HEALTHY | 92.19 | 94.61 | 65.86 | 40.76 | 96.13 | 96.16 | 96.37 |
| LUAD-LUSC | 54.61 | 93.24 | 92.32 | 93.24 | 94.62 | 94.78 | 95.24 |
| LUAD-HEALTHY | 46.16 | 99.25 | 75.78 | 58.59 | 98.96 | 99.10 | 99.25 |
| LUSC-HEALTHY | 51.23 | 99.18 | 73.88 | 56.77 | 99.51 | 99.51 | 99.51 |

### B. Patient-Specific Disparity

Figure 2 shows the patient-specific significant genes reflecting patient-specific heterogeneity or disparity information. Each of the nine Venn diagrams represents three sets of top 100 significant genes derived from three classification problems for a patient or healthy sample, which could be thought of extracting the disparity information using three different types of interrogation. The union of these three sets of genes could be thought of the representation of complete heterogeneity exists in a patient or sample. The top row shows the Venn diagrams of three LUAD patients, first one with the minimum number common genes, third one with the maximum number of common genes and the middle one is an intermediate sample. In similar way, the middle and bottom rows show the samples from LUSC and HEALTY cohorts. The ranges of common genes within the samples of LUAD, LUSC and HEALTHY cohorts are 0 to 17, 0 to 34 and 16 to 40, respectively. The fewer common genes mean the samples are more heterogeneous, which is reflected in LUAD and LUSC cohorts compared to healthy samples, as expected.

**Fig. 2.**
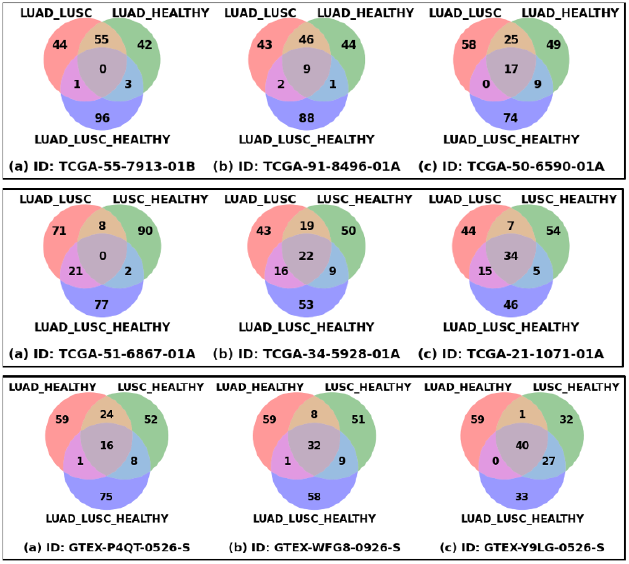
Patient-specific significant genes from three classification problems. Venn diagrams of three LUAD samples (Top Row), three LUSC samples (Middle row), three HEALTHY samples (Bottom Row).

### C. Race and Sex-Specific Correct Prediction

Table IV shows the sub-cohort or race and sex-specific correct prediction for LUAD and LUSC cohorts. For example, actual number of samples in LUAD-AAM sub-cohort is 20, of which at least 18 samples were correctly predicted in all three classification problems. This means that in some classification, the correctly predicted number could be more than 18, and thus, 90% is the minimum of three prediction accuracies. Similarly, LUAD-AAF has a minimum accuracy of 96%.

**TABLE IV.** Race and Sex-Specific Prediction.

| Sub-Cohort | LUAD Cohort |  |  | LUSC Cohort |  |  |
| --- | --- | --- | --- | --- | --- | --- |
|  | Actual | Correct Prediction | Accuracy | Actual | Correct Prediction | Accuracy |
| AAM | 20 | 18 | 90% | 16 | 14 | 88% |
| EAM | 136 | 129 | 95% | 195 | 182 | 93% |
| AAF | 27 | 26 | 96% | 12 | 12 | 100% |
| EAF | 173 | 161 | 93% | 72 | 63 | 88% |
| Total | 356 | 334 | 94% | 295 | 271 | 92% |

### D. Race and Sex-Specific Disparity

Figure 3 shows the Venn diagrams of significant gene sets corresponding to two race (AA and EA) and two sex (M and F) specific sub-cohorts of LUAD (top row) and LUSC (bottom row) cohorts. For example, 984 (15 + 969) significant genes for LUAD-AAM sub-cohort were derived from the union of three sets of 100 significant genes for each of 18 correctly predicted samples (Table III). Similarly, 1,244 (969 + 275) significant genes for LUAD-EAM were derived from the union of significant gene sets for 129 correctly predicted samples.

**Fig. 3.**
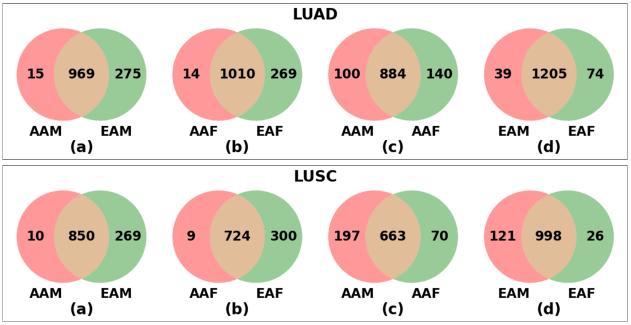
Race and Sex-Specific Significant Gene Sets

Each Venn diagram produced three sets of genes, namely one common set and two sub-cohort-specific sets. For example, Venn diagram (a) of LUAD cohort produced a common set of 969 genes and AAM- and EAM-specific sets of 15 and 275 genes, respectively. The AAM- and EAM-specific gene sets represent the health disparity information between AAMs and EAMs and the common set represents the common behavior/characteristics of LUAD-AAM and LUAD-EAM. Similarly, LUSC-AAM and LUSC-EAM specific 10 and 269 genes represent the lung cancer disparity information for LUSC-AAM and LUSC-EAM, respectively. It is clear from Figure 3 that health disparity exists between AAM and EAM, AAF and EAF, AAM and AAF, and EAM and EAF for both LUAD and LUSC types of lung cancer. This discovery underscores the significance of the proposed approach of delineating lung cancer health disparity, and thus, the disparity related discovered gene sets deserve further validation via wet lab experiment.

### E. Funtional Analysis of Race and Sex-Specific Disparity

The previous section outlined that for the LUAD cohort, the common set of 969 genes represent the common characteristics of AAMs and EAMs and the smaller sets consisting of 15 and 275 genes carry the disparity information for AAMs and EAMs, respectively. To delineate the disparity at functional level, we conducted functional enrichment analyses for two sets of genes reflecting the whole behavior of AAM (15 + 969 = 984 genes) and EAM (275 + 984 = 1,244 genes) cohorts. Figure 4 summarizes the functional enrichment analysis for LUAD-AAM and LUAD-EAM cohorts. Two bar plots show the top 10 significantly enriched GO terms for AAM and EAM cohort of which nine terms are common. The Venn diagram shows that the AAM and EAM cohorts are enriched in 103 and 106 GO terms, respectively. 86 GO terms are common between two cohorts reflecting the common characteristics, whereas 17 and 20 GO terms are solely specific to AAM and EAM cohorts, reflecting the disparity information in LUAD type of lung cancer between AAMs and EAMs.

**Fig. 4.**
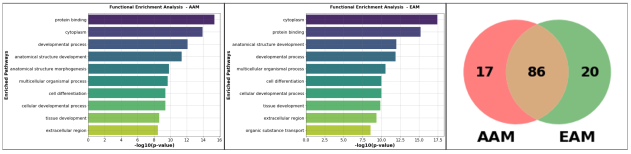
Functional Enrichment Analysis of LUAD-AAM and LUAD-EAM cohorts.

### F. Recovery of Disparity Information: Multiple-Class vs. Single-Class Analysis

Figure 5 compares the performance of multiple-classification approach with single-classification in discovering health disparity in lung cancer between race and sex-specific cohorts. Top row shows the Venn diagrams of the significantly enriched GO terms between two cohorts of AAM vs EAM, AAF vs. EAF, AAM vs. AAF, and EAM vs. EAF using multiple-classification approach and the other three rows show the same using single-classification. In general, the multiple-classification technique provides more pathways enriched for both cohorts compared to single-classification. This means that multiple-classification technique provides more robust information about the disparity in lung cancer.

**Fig. 5.**
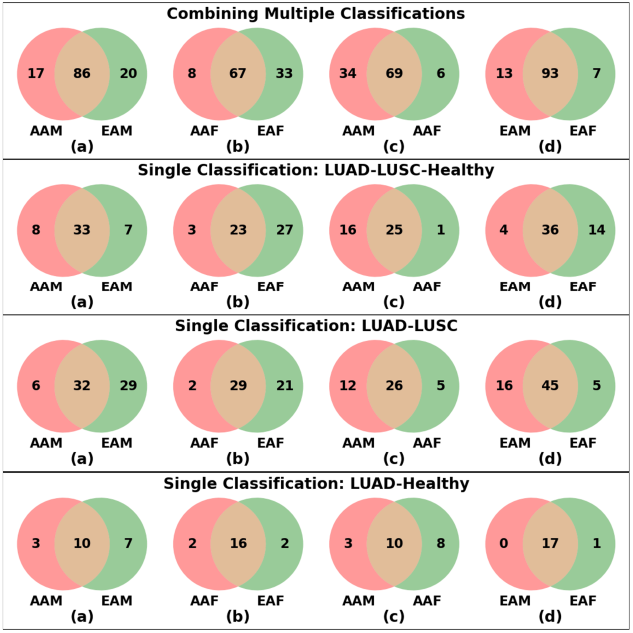
Comparison of Pathway-Level Disparity: Multiple-Classification vs. Single-Classification. Venn Diagrams of Significant GO terms Between Two Cohorts of Race and Sex Combinations.

## IV. Conclusions

This study developed a computational framework to retrieve the disparity information that exists in lung cancers between race and sex-specific cohorts. The proposed approach of interrogating a patient via multiple classifications generates more robust information about disparity compared to interrogation via a single classification. In general, each real-world dataset is highly imbalanced in terms of race, which makes it difficult to retrieve race-specific information. Instead of setting up a classification task based on race labels, we designed it based on disease conditions, which is innovative, and we are the first group to adopt this strategy.

The proposed study successfully discovers the sets of genes and enriched pathways related to health disparity in lung cancer, which deserves to be validated by wet lab experiments. Moreover, the proposed approach can be adapted to discover race- and sex-specific disparity in other cancers and diseases.

## Acknowledgment *(Heading 5)*

This work was supported by the NIH/NHGRI 1UG3HG013615-01, NIH/NCI 1R21CA290324-01 and the State of Florida Biomedical Research Program, Bankhead Coley Research Infrastructure grant number 23B16. The content is solely the responsibility of the authors and does not necessarily represent the official views of the funding agencies.

## References

[1] R. L. Siegel Mph et al., “Cancer statistics, 2023,” CA. Cancer J. Clin., vol. 73, no. 1, pp. 17–48, Jan. 2023.

[2] F. Islami et al., “Proportion and number of cancer cases and deaths attributable to potentially modifiable risk factors in the United States,” CA. Cancer J. Clin., vol. 68, no. 1, pp. 31–54, Jan. 2018.

[3] K. A. Mitchell, A. Zingone, L. Toulabi, J. Boeckelman, and B. M. Ryan, “Comparative transcriptome profiling reveals coding and noncoding RNA differences in NSCLC from African Americans and European Americans,” Clin. Cancer Res., vol. 23, no. 23, pp. 7412–7425, Dec. 2017.

[4] J. N. Sampson et al., “Analysis of heritability and shared heritability based on genome-wide association studies for thirteen cancer types,” J. Natl. Cancer Inst., vol. 107, no. 12, 2015.

[5] K. A. Zanetti et al., “Genome-wide association study confirms lung cancer susceptibility loci on chromosomes 5p15 and 15q25 in an African-American population,” Lung Cancer, vol. 98, pp. 33–42, Aug. 2016.

[6] J. McKay, R. Hung, Y. Han, … X. Z.-N., and U. 2017, “Large-scale association analysis identifies new lung cancer susceptibility loci and heterogeneity in genetic susceptibility across histological subtypes,” Nat. Genet., vol. 49, pp. 1126–1132, 2017.

[7] A. Al Mamun, M. Sobhan, R. B. Tanvir, C. J. Dimitroff, and A. M. Mondal, “Deep Learning to Discover Cancer Glycome Genes Signifying the Origins of Cancer,” in 2020 IEEE International Conference on Bioinformatics and Biomedicine, BIBM, Dec. 2020, pp. 2425–2431.

[8] A. Al Mamun et al., “Multi-Run Concrete Autoencoder to Identify Prognostic lncRNAs for 12 Cancers,” Int. J. Mol. Sci., vol. 22, no. 21, p. 11919, Nov. 2021.

[9] M. Sobhan, A. Al Mamun, R. B. Tanvir, M. J. Alfonso, P. Valle, and A. M. Mondal, “Deep Learning to Discover Genomic Signatures for Racial Disparity in Lung Cancer,” in 2020 IEEE International Conference on Bioinformatics and Biomedicine BIBM, Dec. 2020, pp. 2990–2992.

[10] S. M. Lundberg and S. I. Lee, “A Unified Approach to Interpreting Model Predictions,” Adv. Neural Inf. Process. Syst., vol. 2017-Decem, pp. 4766–4775, May 2017.

[11] M. Sobhan and A. M. Mondal, “Explainable Machine Learning to Identify Patient-specific Biomarkers for Lung Cancer,” in 2022 IEEE International Conference on Bioinformatics and Biomedicine (BIBM), 2022, pp. 3152–3159.

[12] M. Verma, “Personalized Medicine and Cancer,” J. Pers. Med., vol. 2, pp. 1–14, 2012.

[13] Using Population Descriptors in Genetics and Genomics Research: A New Framework for an Evolving Field. National Academies Press, 2023.

[14] T. M. Grzywa, W. Paskal, and P. K. Włodarski, “Intratumor and Intertumor Heterogeneity in Melanoma,” Transl. Oncol., vol. 10, no. 6, pp. 956–975, Dec. 2017.

[15] Q. Wang et al., “Unifying cancer and normal RNA sequencing data from different sources,” Sci. Data 2018 51, vol. 5, no. 1, pp. 1–8, Apr. 2018.

[16] G. F. Gao et al., “Before and After: Comparison of Legacy and Harmonized TCGA Genomic Data Commons’ Data,” Cell Syst., vol. 9, no. 1, pp. 24-34.e10, Jul. 2019.

[17] J. Lonsdale et al., “The Genotype-Tissue Expression (GTEx) project,” Nat. Genet. 2013 456, vol. 45, no. 6, pp. 580–585, May 2013.

[18] M. Sobhan and A. M. Mondal, “Evaluating SHAP’s Robustness in Precision Medicine: Effect of Filtering and Normalization,” in Proceedings - 2023 IEEE International Conference on Bioinformatics and Biomedicine, BIBM, 2023, pp. 3157–3164.

